# ASTRA enables zero-shot super-resolution of spatial transcriptomics across capture geometries

**DOI:** 10.64898/2026.09.30.755663

**Authors:** Jiachen Ye, Qinghua Cui

**Affiliations:** Department of Biomedical Informatics, State Key Laboratory of Vascular Homeostasis and Remodeling, School of Basic Medical Sciences, Peking University, 38 Xueyuan Rd, Beijing, 100191, China; School of Sports Medicine, Wuhan Sports University, No. 461 Luoyu Rd., Wuhan 430079, Hubei Province, China

**Keywords:** Spatial transcriptomics, Super-resolution, Pretrained model, Zero-shot inference, Capture geometry, Cohort analysis

## Abstract

Spatial transcriptomic super-resolution recovers tissue organization obscured by coarse sampling, but fitting a model to each section greatly increases computational cost and reduces practicality. We introduce ASTRA, a cross-platform pretrained model that reconstructs expression from histology and measured RNA by zero-shot inference. Training with arbitrarily shaped count-bearing regions, termed parents, enables one frozen model to accommodate dense bins and separated spots. Across six held-out Visium HD sections, ASTRA achieved higher correlations under dense-bin and pseudo-Visium sampling, lower dense-bin errors and substantially shorter processing times than existing methods. It improved colorectal tissue segmentation and tumor-boundary recovery. In real colorectal Visium, reconstruction spatially resolved a stromal transcriptional state supported by measured RNA. The same checkpoint supported regional and nucleus-associated analyses across 43 invasive lobular carcinoma sections and within-spot interpretation across 92 triple-negative breast cancer sections. ASTRA establishes a reusable framework for practical, cohort-scale spatial transcriptomic super-resolution across platforms.

## Main

Spatial transcriptomics connects molecular states to the tissue structures in which they operate^1^. In tumors, malignant nests, stromal territories and immune neighborhoods meet at boundaries that rarely coincide with capture units^2^. A single spot can therefore mix biologically distinct compartments, concealing how their molecular programs are arranged. Even high-definition measurements are often aggregated to increase molecular support, trading spatial detail for more robust expression profiles^2^. Recovering organization below these observation scales would extend both the biological utility of existing datasets and the reach of cohort studies.

Computational super-resolution uses molecular, spatial and histological information to recover this lost detail. BayesSpace uses Bayesian spatial modeling to resolve subspot variation^3^, whereas STAGE fits a spatial location-supervised autoencoder to generate expression at unmeasured positions^4^. XFuse jointly models histology and transcript counts through a deep generative model^5^. TESLA uses spatially and histologically weighted interpolation^6^, and Thor propagates expression over a morphology-informed cell graph by anti-shrinking Markov diffusion^7^; their expression-enhancement stages do not require model training. iStar combines hierarchical image features with expression prediction^8^, FineST aligns image and RNA representations through contrastive learning^9^, and scstGCN integrates morphological and spatial features through graph convolution^10^. These neural reconstruction methods use measured spot expression as weak supervision for fine-scale predictions. The recent FineST benchmark showed that super-resolution with target-section fitting can take hours for a single section, as reported for FineST and iStar on colorectal tissue^9^. SpotZoomer uses reference-guided transfer learning, leveraging high-resolution ST data—typically from biologically related tissues—as a teacher, followed by teacher–target domain alignment and spot-level weakly supervised fine-tuning on the target slice^11^. Its published workflow therefore depends on reference training and target adaptation and does not provide a pretrained reconstruction model for direct zero-shot inference. Fitting a reconstruction model to each section increases the cost of analyzing large cohorts, while reference-guided transfer depends on suitable high-resolution data. Reusing pretrained weights across new sections would eliminate target-section fitting and make large-scale analysis more practical.

Zero-shot reconstruction across assay platforms requires a model to accommodate differences in capture geometry. Visium HD provides contiguous square bins, whereas conventional Visium and earlier spatial transcriptomic platforms measure separated circular spots of different sizes and spacing^1,2^. SRast has demonstrated generalization across platforms using a pretrained flow-matching model coupled to target-specific feature adaptation^12^. However, its released evaluation workflow first aggregates existing measurements into synthetic coarse observations and then reconstructs expression at the original measurement locations^12^. These coarse inputs are formed by grouping measured positions, without explicitly reproducing the capture geometry of a target assay. The resulting benchmarks therefore assess recovery after artificial coarsening, but do not establish super-resolution beyond the native sampling scale or reconstruction across unmeasured inter-spot regions. Extending reconstruction to these gaps is important for recovering continuous tissue organization from sparsely sampled sections. A model for zero-shot reconstruction therefore needs to retain the spatial extent of each measured count while accounting for changes in support shape, area and tissue coverage. Representing observations through their count-bearing regions would provide a common input for dense bins and sparse spots, allowing one pretrained model to serve different assay formats.

Here we introduce ASTRA, Assay-aware Spatial Transcriptomic Reconstruction and Allocation, for zero-shot super-resolution from registered hematoxylin and eosin (H&E) images and coarse RNA. ASTRA uses an arbitrary-parent representation of count-bearing regions and learns from high-definition data aggregated over parents of varied shape and size, allowing the same model to accommodate different sampling patterns. Once pretrained, the same frozen model reconstructs new sections without target-specific training or test-time optimization; optional self-supervised fine-tuning is evaluated separately. Across six held-out sections, ASTRA achieves higher reconstruction correlations and lower dense-bin errors with substantially shorter processing times. In colorectal cancer (CRC), reconstruction improves tissue segmentation and tumor-boundary recovery, and connects cell-associated RNA sources to the spatial organization of a stromal transcriptional state supported by measured expression. We then apply the same checkpoint to 43 invasive lobular carcinoma (ILC) and 92 triple-negative breast cancer (TNBC) sections. These applications establish the practical value of pretrained super-resolution for regional, nucleus-associated and molecular-program analyses across conventional Visium and the larger capture spots of the original Spatial Transcriptomics platform.

## Results

### Arbitrary-parent training enables frozen inference across capture geometries

ASTRA receives H&E, measured counts and parent-support masks, and predicts expression on an 8 µm grid (Fig. 1a). During training, native high-definition measurements are aggregated over randomly generated parents. Learning to reconstruct from these changing supports enables deployment across contiguous bins and separated spots. The source model was fitted on 12 development sections using a fixed 2,000-gene panel, with spatially withheld regions for model selection. Test sections and downstream cohorts were excluded from fitting and selection (Supplementary Methods 1 and 2; Supplementary Table 1).

**Figure 1.**
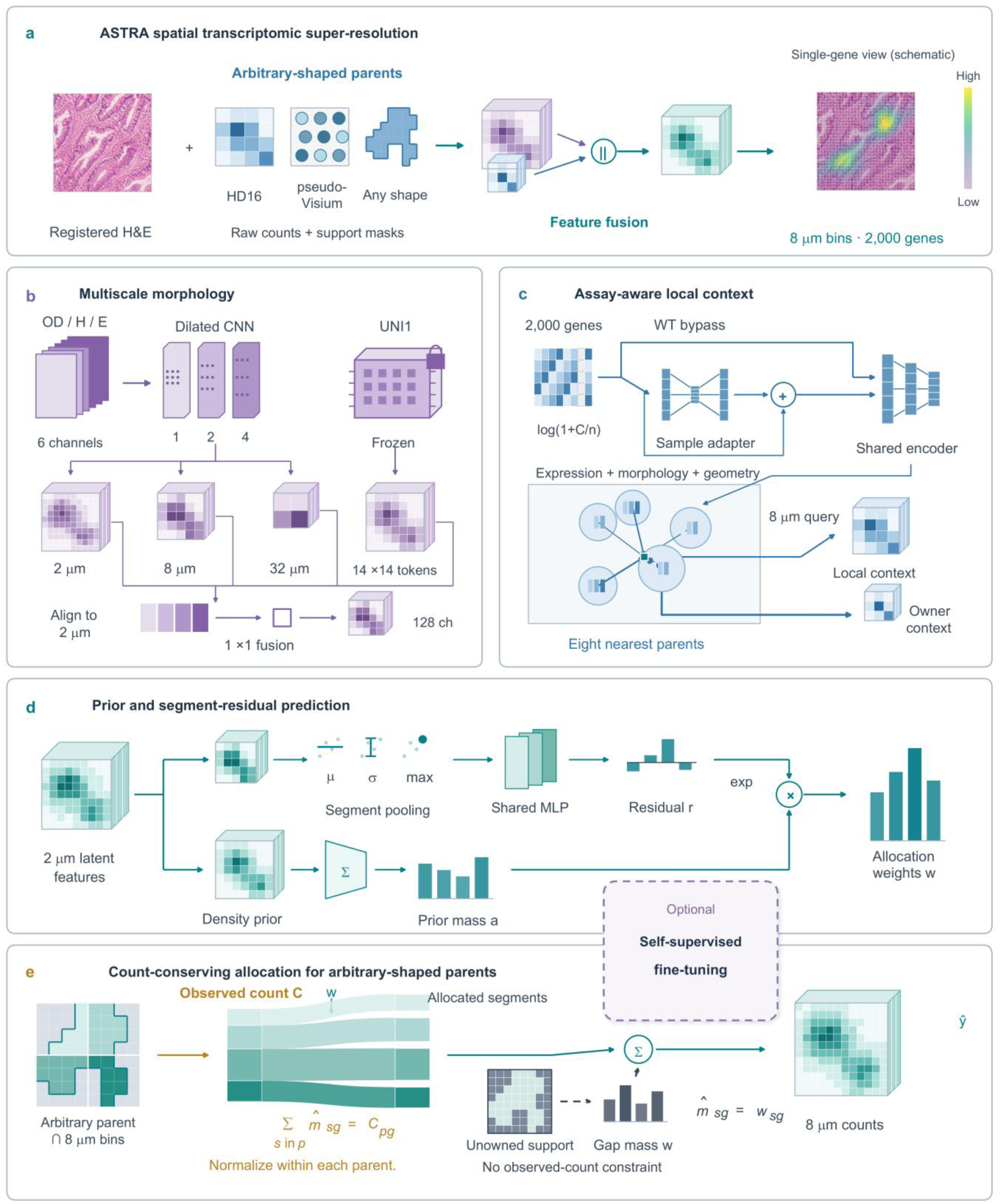
Arbitrary-parent representation supports zero-shot reconstruction across capture geometries. **a**, ASTRA accepts registered H&E, measured counts and support masks and produces an 8 µm grid for 2,000 genes. Square bins, circular spots and arbitrary shapes share the same parent representation. HD16 and pseudo-Visium illustrate the evaluated dense and sparse geometries; source training uses randomly generated parent shapes, areas and coverage. The overlaid gene map and colored feature tensors are schematics, not experimental expression results. Assay identity and gene availability are separate inputs. **b**, Local optical-density (OD), hematoxylin (H) and eosin (E) features are combined with multiscale convolutional branches and frozen UNI spatial tokens. The internal 2 µm grid represents image and support geometry; RNA predictions are exported on the 8 µm grid. **c**, Count-density features pass through a sample adapter for 3′ measurements or a whole-transcriptome (WT) bypass before shared molecular encoding. Parent tokens combine expression, morphology and geometry. Attention reads up to eight nearest valid parents without a fixed radius cutoff; owned cells also receive their parent’s token. Gap density uses local parent context, with a global fallback only when that context is empty. **d**, The upper segment-residual branch and lower density-prior branch yield positive allocation weights w from the product of integrated prior mass a and the exponentiated residual r. Segment pooling summarizes cell features by their mean, standard deviation and maximum. **e**, Normalization within each observed parent allocates its measured gene counts to intersections with 8 µm bins. Segment contributions are then summed into the output grid. Their sum equals each observed parent/gene total, whereas gap mass has no local observed-total constraint. Symbols follow Methods, equations (1)–(4); whole-section overlap averaging is distinct from this field-level constraint. The floating dashed module between d and e denotes optional self-supervised fine-tuning, which merges coarse supports/counts, predicts the original coarse observations and preserves frozen source allocations, selecting a checkpoint on separate coarse regions. This branch uses no target fine-resolution labels and is reported separately as ASTRA fine-tuned. The primary ASTRA results use frozen-weight inference.

Local stain features, multiscale convolutional features and frozen UNI pathology features encode morphology^13^ (Fig. 1b). Molecular encoding combines count density, gene availability and assay identity. Parent tokens integrate expression, morphology and geometry, and provide local molecular context to each query (Fig. 1c). A density-prior branch and a segment-residual branch jointly determine allocation weights (Fig. 1d). Normalization within each parent conserves its measured gene counts when allocating them to intersecting output bins, while learned density estimates supply expression in unmeasured gaps (Fig. 1e). Whole-section predictions combine overlapping fields (Methods).

The primary deployment uses frozen ASTRA weights. Optional self-supervised fine-tuning uses only coarse target observations and is evaluated separately as ASTRA fine-tuned.

### Six-section benchmarks show accurate reconstruction with substantially shorter processing times

We compared ASTRA with FineST, iStar, scstGCN and TESLA on six held-out Visium HD sections: two ovarian, one breast, CRC P2, one colon and one pancreas section. In the dense-bin benchmark, 16 µm observations were reconstructed on an 8 µm grid using the same 2,000 genes and common scoring support (Fig. 2a; Supplementary Method 3; Supplementary Table 2).

**Figure 2.**
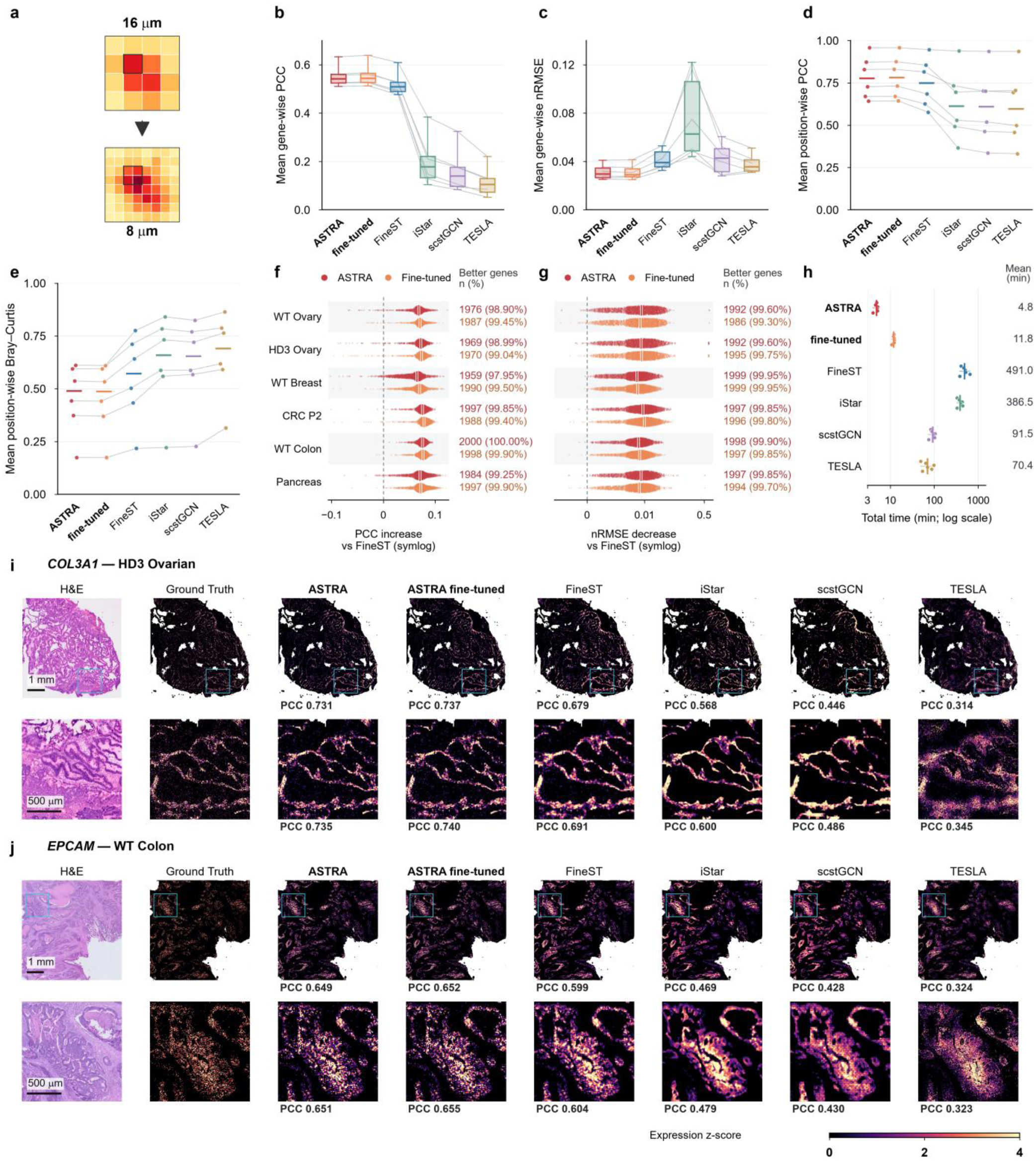
Reconstruction accuracy and recorded runtime across six tissue sections. ASTRA, ASTRA fine-tuned, FineST, iStar, scstGCN and TESLA are compared on 2,000 genes in six sections: WT Ovarian, HD3 Ovarian, WT Breast Ultima, CRC P2, WT Colon and WT Pancreas. **a**, Schematic of 16-to-8 µm reconstruction. **b,c**, Section means of gene-wise PCC and nRMSE. Lines pair sections; boxes show interquartile ranges, center lines medians and whiskers the full range. PCC uses count-scale expression and 1,998/1,989/2,000/2,000/2,000/1,999 defined genes, respectively; nRMSE uses independent per-gene min–max scaling and all 2,000 genes. **d,e**, Section means of position-wise PCC and Bray–Curtis dissimilarity. Points are sections; lines pair them and colored marks indicate medians. **f,g**, Gene-wise improvement relative to FineST, the highest-PCC external method in every section: ASTRA minus FineST PCC (f), and FineST minus ASTRA nRMSE (g). Positive values favor ASTRA. All finite pairs are shown; marks indicate medians and annotations report strict-improvement counts and percentages. Vertical displacement separates genes. Symmetric-log axes are linear within ±0.02 PCC and ±0.003 nRMSE. **h**, Recorded stage totals on a logarithmic axis. Points denote sections, horizontal lines their range, and vertical marks and numbers their arithmetic mean. Totals sum recorded preparation, target fitting where applicable and inference stages, excluding source pretraining. **i,j**, Whole-section and enlarged *COL3A1* maps in HD3 Ovarian and *EPCAM* maps in WT Colon. Cyan boxes mark 1.536-mm ROIs. Each map uses whole-support standardization and a shared z-score range of 0–4 without ROI rescaling. PCC uses unclipped expression, including zeros. Scale bars, 1 mm and 500 µm. Examples were selected post hoc for visualization. Scoring support, display transformations and timing definitions are given in Supplementary Method 3 and Supplementary Table 2.

Frozen and fine-tuned ASTRA achieved a higher mean gene-wise Pearson correlation coefficient (PCC) and a lower normalized root mean squared error (nRMSE) than every external method in each section (Fig. 2b,c). Unweighted mean PCC across sections was 0.5520 for frozen ASTRA and 0.5554 after fine-tuning, versus 0.5201 for FineST, the strongest external comparator. Corresponding nRMSE values were 0.03124, 0.03119 and 0.04152. Relative to FineST, frozen ASTRA improved PCC for 97.95–100% of evaluable genes and nRMSE for 99.60–99.95% across sections (Fig. 2f,g; Supplementary Figs. 1–4). Fine-tuning increased mean PCC in all six sections, but decreased nRMSE in four.

ASTRA also improved position-wise expression profiles. Across section means, frozen ASTRA increased position-wise PCC from 0.7504 for FineST to 0.7829 and reduced Bray–Curtis dissimilarity from 0.5469 to 0.4553 (Fig. 2d,e). Both ASTRA variants led all external methods on these measures in every section. *COL3A1* in HD3 Ovarian and *EPCAM* in WT Colon illustrate the spatial reconstructions (Fig. 2i,j); marker comparisons and twelve further examples cover all six sections (Supplementary Figs. 5–13). Recorded processing time averaged 4.79 min per section for frozen ASTRA and 11.84 min with fine-tuning, compared with 70.40–490.97 min for external methods (Fig. 2h; Supplementary Fig. 14). Frozen ASTRA therefore reduced mean processing time by 93.2–99.0%. These totals include preparation, target fitting where applicable and inference, and exclude source pretraining.

The same checkpoint reconstructed the six sections from pseudo-Visium observations with a 55 µm spot diameter and 100 µm spacing. This benchmark additionally included SpotZoomer, supplied with a biologically related high-resolution training section for each target (Supplementary Fig. 15; Methods). Both ASTRA variants achieved higher gene-wise and position-wise PCC than every external method in each section. Mean gene-wise PCC was 0.1663 for frozen ASTRA and 0.1698 after fine-tuning, versus 0.1394 for iStar, the highest external mean, and 0.0982 for SpotZoomer. Error rankings differed: scstGCN had the lowest nRMSE in every section, averaging 0.04426 versus 0.05514 for frozen ASTRA, and lower Bray–Curtis dissimilarity in HD3 Ovarian and CRC P2. Frozen ASTRA required a mean recorded processing time of 4.84 min, versus 11.23–58.90 min for external methods, including 47.94 min for SpotZoomer’s reference–target workflow. Section-specific values are provided in Supplementary Fig. 16. The same pretrained checkpoint therefore retained its correlation and runtime advantages under sparse capture geometry.

### ASTRA improves colorectal tissue segmentation and tumor-boundary recovery

We next tested whether expression reconstruction improved tissue segmentation and tumor-boundary prediction in colorectal P2 pseudo-Visium data. A shared classifier fitted to native 8 µm expression and published pathology annotations^14^ mapped native and reconstructed profiles to four tissue compartments; 3,806 simulated spots supplied the coarse inputs (Fig. 3; Supplementary Method 4).

**Figure 3.**
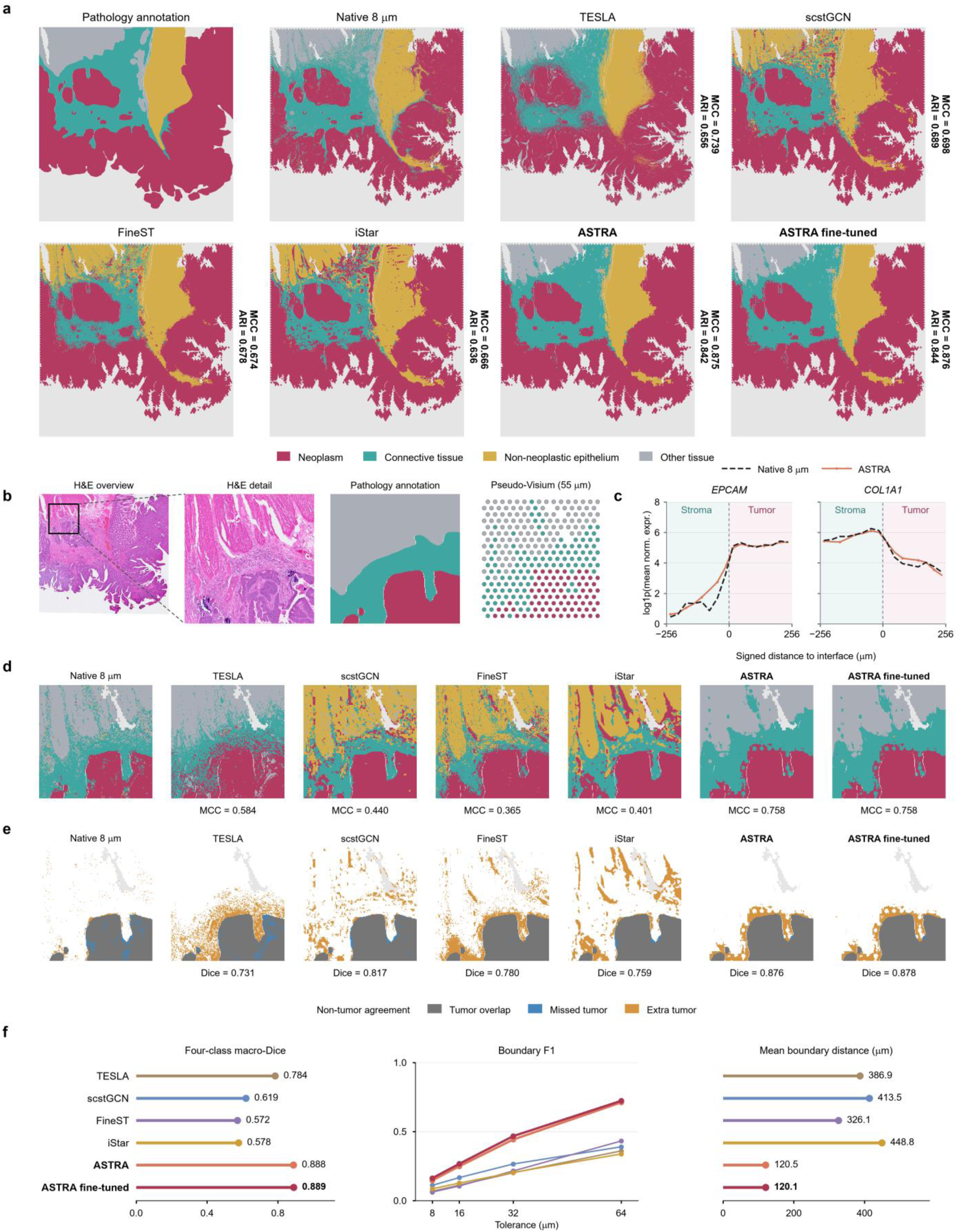
Improved colorectal tissue segmentation and tumor-boundary recovery. **a**, Whole-section pathology, native-derived partition and six reconstructed tissue maps on the same capture grid. A fixed multinomial classifier fitted to native expression and annotations assigns four tissue classes. Method annotations report pathology MCC and adjusted Rand index over 493,080 common annotated bins. Gray marks unlabeled or excluded support; native scores are omitted. **b**, H&E overview, 1.536 × 1.536 mm enlargement, transferred pathology and pseudo-Visium inputs. Spot colors use the same classifier and tissue key. **c**, Native and ASTRA *EPCAM* and *COL1A1* profiles across the annotated tumor–stroma interface. The same 16,415 bins within 256 µm of the interface enter sixteen 32 µm bands; distance is negative in stroma and positive in tumor. Counts are normalized per bin over 2,000 genes, averaged within bands and then log1p transformed. Both genes share the scale used in Supplementary Fig. 20. Lines join unsmoothed band means; background shading identifies tumor and stroma. **d,e**, Local tissue assignments and tumor-overlap maps in b. MCC uses 35,777 annotated bins; local tumor Dice is reported separately. Dark gray denotes overlap, blue missed tumor, orange additional predicted tumor and white non-tumor agreement. Native maps are fitted references without displayed scores. **f**, Whole-section macro-Dice, boundary F1 at 8/16/32/64 µm tolerances and mean symmetric boundary distance. Macro-Dice equally weights four classes. Boundary metrics pool all evaluable boundaries, including predictions outside the displayed window, against 2,010 reference pixels after support-edge exclusion. Distance pools nearest-neighbor distances in both directions. Scores summarize this section; matching tolerances specify allowable boundary displacement. Counts use Space Ranger 4.0.1; annotations were transferred from the Space Ranger 3.0 coordinates through the common H&E. Full definitions and regional sensitivities are in Supplementary Method 4, Supplementary Figs. 17–20 and Supplementary Table 3.

Across 493,080 common annotated bins, frozen ASTRA achieved a Matthews correlation coefficient (MCC) of 0.875 and an adjusted Rand index of 0.842 against pathology, exceeding the external methods’ MCCs of 0.666–0.739 and adjusted Rand indices of 0.636–0.689 (Fig. 3a; Supplementary Table 3). Tumor Dice reached 0.9658, compared with 0.904–0.940 for the comparators. Class-balanced macro-Dice was 0.888, versus 0.572–0.784, with gains in every tissue compartment (Fig. 3f; Supplementary Fig. 17).

A 1.536-mm field connects these segmentation results to local histology and expression (Fig. 3b). *EPCAM* and *COL1A1* profiles across the pathology-defined tumor–stroma interface show how reconstructed epithelial and matrix-associated RNA vary across the boundary (Fig. 3c). Local tissue assignments and tumor-overlap maps distinguish recovered compartments from missed and additional predicted tumor (Fig. 3d,e).

Across the entire evaluable section, frozen ASTRA achieved boundary F1 scores of 0.146, 0.249, 0.442 and 0.711 at 8, 16, 32 and 64 µm tolerances, exceeding every comparator at each tolerance (Fig. 3f). Mean symmetric boundary distance decreased to 120.53 µm, versus 326.07–448.83 µm for external methods. Fine-tuning further increased boundary F1 to 0.162, 0.266, 0.467 and 0.723. Capture-domain analyses, paired regional effects and comparator expression profiles provide complementary views of tissue recovery and boundary localization (Supplementary Figs. 18–20).

### ASTRA resolves the spatial organization of a stromal transcriptional state in real Visium

We then examined what reconstructed spatial information could reveal in experimentally acquired colorectal Visium. The P2 section contained 4,269 measured spots and yielded 589,267 reconstructed positions. Using a single reference derived from four non-P2 donors, we projected Native and ASTRA expression onto cell-associated RNA sources^2^. The resulting maps depicted finer local arrangements of epithelial-associated, fibroblastic, mural and endothelial RNA (Fig. 4a). Reference qualification, synthetic-mixture checks and omission of program genes supported their interpretation as estimated RNA contributions (Supplementary Fig. 21a–c).

**Figure 4.**
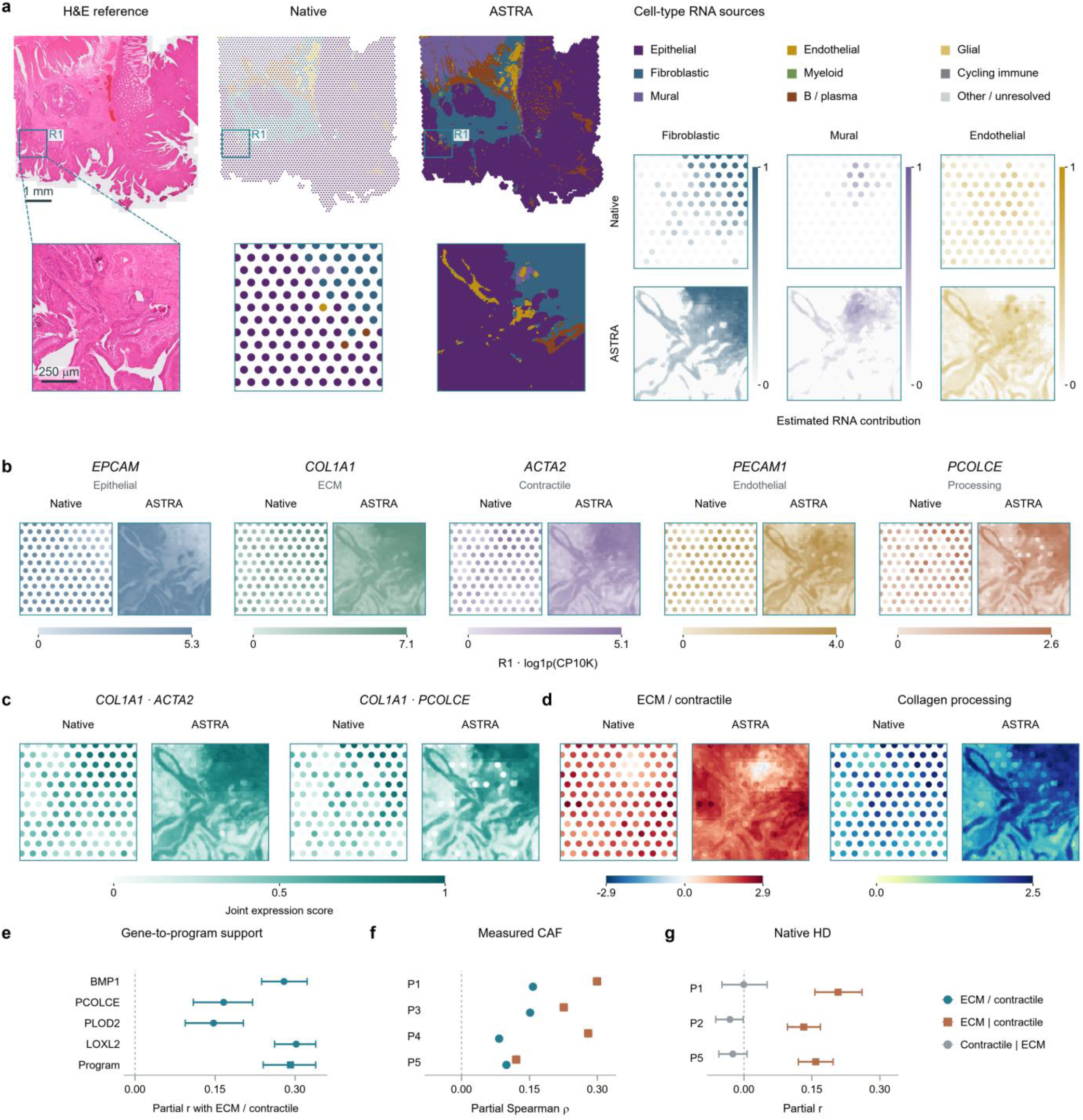
Cell-associated RNA sources and molecular programs in real colorectal Visium. **a**, H&E, Native and ASTRA source maps of P2 CRC (4,269 spots; 589,267 reconstructed 8-µm positions). Weighted non-negative least squares (NNLS) uses four non-P2 reference donors. Categorical colors indicate the dominant estimated RNA contribution; gray denotes unresolved assignments. Paired continuous maps show fibroblastic, mural and endothelial contributions. R1 is the same 1,024-µm region throughout a–d. Scale bars, 1 mm and 250 µm. **b**, Representative genes, expressed as log1p(CP10K) over the shared 2,000-gene panel. Each gene has one shared Native/ASTRA color range. **c**, Joint expression score, J = min(u, v), where u and v are relative gene intensities clipped to 0–1 using the shared limits in b. Higher scores indicate jointly high relative gene expression. **d**, Natural-log ECM/contractile mean-count ratio and mean log1p(CP10K) for *BMP1*, *PCOLCE*, *PLOD2* and *LOXL2*. **e**, Adjusted within-spot associations of ECM/contractile balance with each processing gene and their program. Points are mean partial Pearson correlations across 48 spatial blocks; bars are 95% block-bootstrap intervals. **f**, Partial Spearman correlations in measured CAFs from P1/P3/P4/P5 (630/389/1,197/950 eligible cells). Symbols distinguish the balance endpoint from ECM conditional on contractile expression. **g**, Conditional ECM and contractile associations with processing in native P1/P2/P5 HD (23/39/36 spatial blocks), shown as block means and 95% bootstrap intervals. Vertical bars in the shared key denote conditioning. Spatial blocks are 1,024 µm; intervals use 1,999 resamples. CAF and HD panels use complementary measurements from the same study. Source maps show estimated RNA contributions. Program genes, qualification and covariates are detailed in Supplementary Method 5 and Supplementary Table 4.

Within the same fixed region, *EPCAM* and *PECAM1* connected epithelial and endothelial source patterns to individual genes, whereas *COL1A1*, *ACTA2* and *PCOLCE* linked local expression to matrix production, contractility and collagen processing (Fig. 4b). A joint expression score, defined by the lower of two shared-scale relative gene intensities, displayed where *COL1A1–ACTA2* and *COL1A1–PCOLCE* signals were jointly high (Fig. 4c). The pairs were positively correlated across Native spots, and ASTRA resolved where their joint expression was concentrated within tissue territories.

Multigene maps and analyses across methods and spatial supports linked these local patterns to broader stromal transcription (Fig. 4d; Supplementary Fig. 21d,f–h; Supplementary Table 4). The within-spot association between extracellular-matrix (ECM)-to-contractile balance and collagen-processing expression remained positive after adjustment for depth, tissue and vascular backgrounds, nuclear density and *NOTCH3* (partial r = 0.292; 95% spatial-block bootstrap interval, 0.240–0.339). All four processing genes showed positive associations (Fig. 4e).

Measured RNA supported the link between ECM production and collagen-processing transcription. Cancer-associated fibroblasts (CAFs) from four non-P2 donors showed positive associations between ECM-to-contractile balance and collagen processing, including an ECM contribution conditional on contractile expression (Fig. 4f). An eight-gene set outside the ASTRA panel supplied additional measured-expression support (Supplementary Fig. 21e). Native HD measurements from three donors likewise supported the conditional ECM association (partial r = 0.132–0.208; Fig. 4g), with regional tumor-proximity analyses providing context (Supplementary Fig. 21i). These findings support a stromal transcriptional state coupling ECM production to collagen processing, consistent with matrix-remodeling CAF phenotypes in CRC^15,16^. ASTRA spatially resolved this ECM–processing relationship in real Visium, connecting measured stromal transcription to local tissue organization.

### Zero-shot reconstruction extends regional and nucleus-associated analysis across an ILC cohort

Zero-shot inference allowed ASTRA to extend regional and nucleus-associated analysis across a patient cohort without fitting a reconstruction model for each patient. The same frozen checkpoint reconstructed all 43 primary invasive lobular carcinoma (ILC) sections in the Visium cohort of Serra et al.^17^ (Fig. 5; Supplementary Table 5). The cohort comprised 71,997 quality-controlled spots and yielded 9.96 million predicted positions. The inference queue completed in 60.76 min using four NVIDIA A100 GPUs (80 GB each), excluding registration and preflight checks.

**Figure 5.**
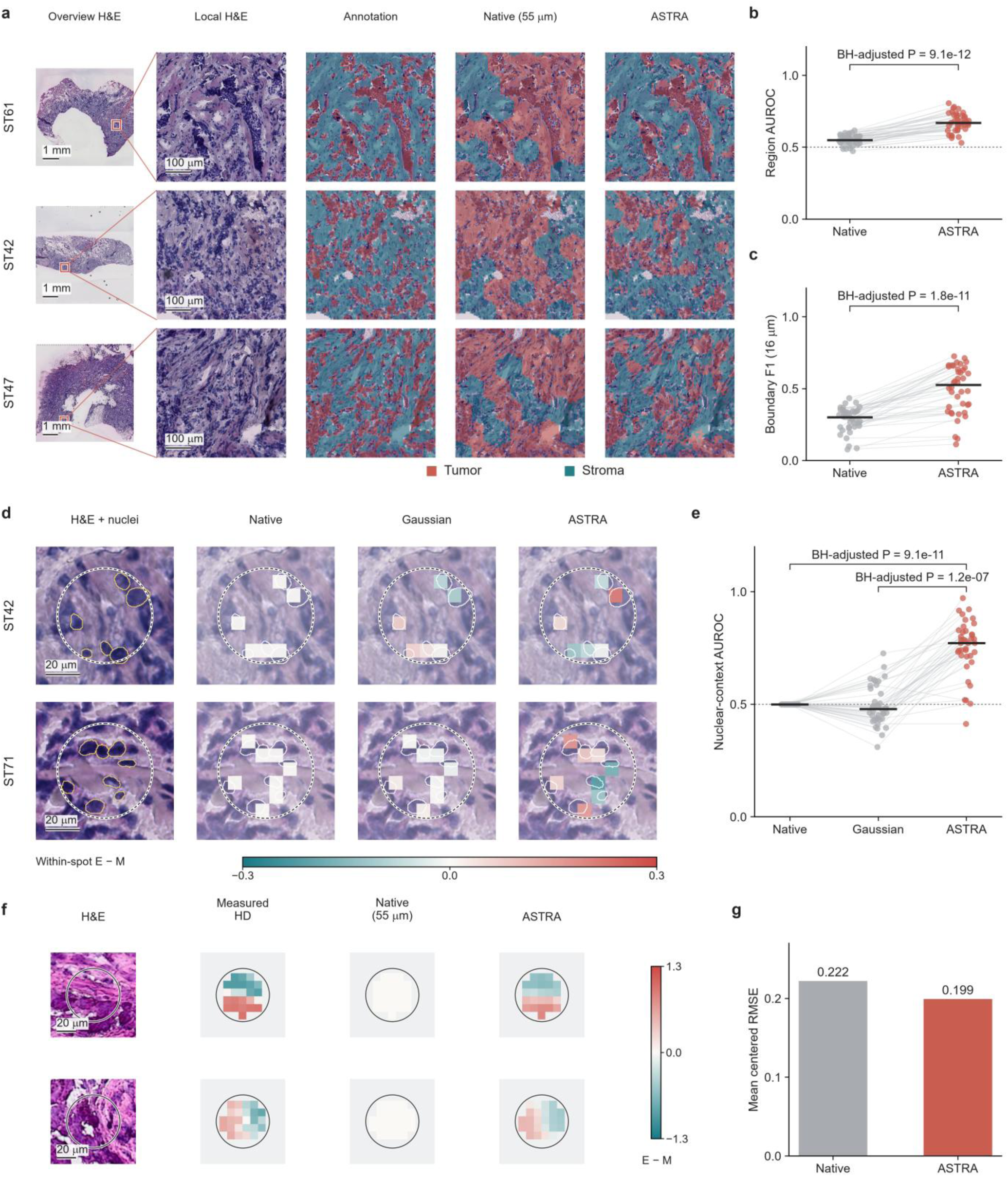
Zero-shot cohort reconstruction supports regional and nucleus-associated RNA analysis in ILC. **a**, H&E, pathology and RNA-based regional classifications. **b,c**, Regional AUROC and 16-µm boundary F1 (40 patients; 116 ROIs). **d**, Nuclear neighborhoods; outlines mark retained nuclei. **e**, Nuclear-context AUROC (39 patients). **b,c,e**, Paired patient points and median bars; two-sided paired Wilcoxon tests, Benjamini–Hochberg (BH)-adjusted P values (b/e: five comparisons including Supplementary Fig. 22b,c; c: ten comparisons). **f**, Two simulated 55-µm spots from the breast HD benchmark section; views span 88 µm. E−M denotes epithelial (cancer + normal) minus mesenchymal (CAF + perivascular-like) RNA contributions, centered within each spot separately by method; red/teal indicate positive/negative deviations. **g**, Mean centered E−M RMSE across 3,789 spots and 87,742 bins. Scale bars: 1 mm, 100 µm and 20 µm. Nuclear contexts are assigned from H&E morphology.

Fixed pathological and spatial-support criteria identified 116 tumor–stroma interface regions in 40 patients. A patient-held-out classifier trained on original-resolution RNA from other patients was applied unchanged to Native and ASTRA expression (Fig. 5a). Across patients, the median regional area under the receiver operating characteristic curve (AUROC) increased from 0.550 to 0.670, and median boundary F1 at 16 µm increased from 0.301 to 0.526; both improved in every evaluable patient (Fig. 5b,c). Macro-Dice increased from 0.487 to 0.537, although overlap gains were concentrated in stroma (Supplementary Fig. 22a–c).

An external breast-cancer single-cell reference^18^ provided a common RNA-source interpretation across patients. Within mixed spots, epithelial-minus-mesenchymal scores distinguished H&E-derived tumor and stromal nuclear contexts with a patient-median AUROC of 0.772 across 39 patients, compared with 0.479 for Gaussian interpolation and 0.5 for spatially constant Native expression (Fig. 5d,e). In a breast HD calibration with simulated 55 µm spots, ASTRA reduced the RMSE of within-spot-centered RNA-source scores by 10.3% relative to Native allocation (Fig. 5f,g). ASTRA thus resolved epithelial and mesenchymal RNA patterns within mixed spots in relation to local nuclear morphology.

Fine-tuning on a small subset of patients further reduced measured-RNA error in a separate adaptation experiment. With five patients for adaptation, three for validation and 35 for testing, the selected model reduced mean Poisson deviance per unique molecular identifier (UMI) by 3.2%, from 0.6882 to 0.6659, when recovering measured spot counts from merged inputs (Supplementary Fig. 22d). Error decreased in all 35 test patients.

### Arbitrary-parent representation enables within-spot analysis of a TNBC cohort on a coarser platform

ASTRA’s arbitrary-parent design enables the same pretrained model to reconstruct spatial transcriptomic data acquired with different sampling geometries. We therefore applied the same frozen checkpoint to the 92-patient TNBC cohort of Wang et al.^19^, acquired with the original Spatial Transcriptomics platform^1^. The platform’s spots are 100 µm in diameter and spaced 150 µm apart, pooling RNA over larger tissue areas than conventional Visium spots. ASTRA reconstructed one primary-tumor section per patient without patient-specific fitting, converting 93,901 quality-controlled spots into 34.91 million positions on an 8 µm grid. We focused on the spatial organization of RNA within these large capture spots.

We first examined a tertiary lymphoid structure (TLS) annotated by the original study authors in TNBC33. Native measurements provided one TLS-associated score per capture spot. ASTRA resolved higher scores inside the annotated TLS and lower scores in adjacent tissue, including within two spots crossed by its boundary (Fig. 6a).

**Figure 6.**
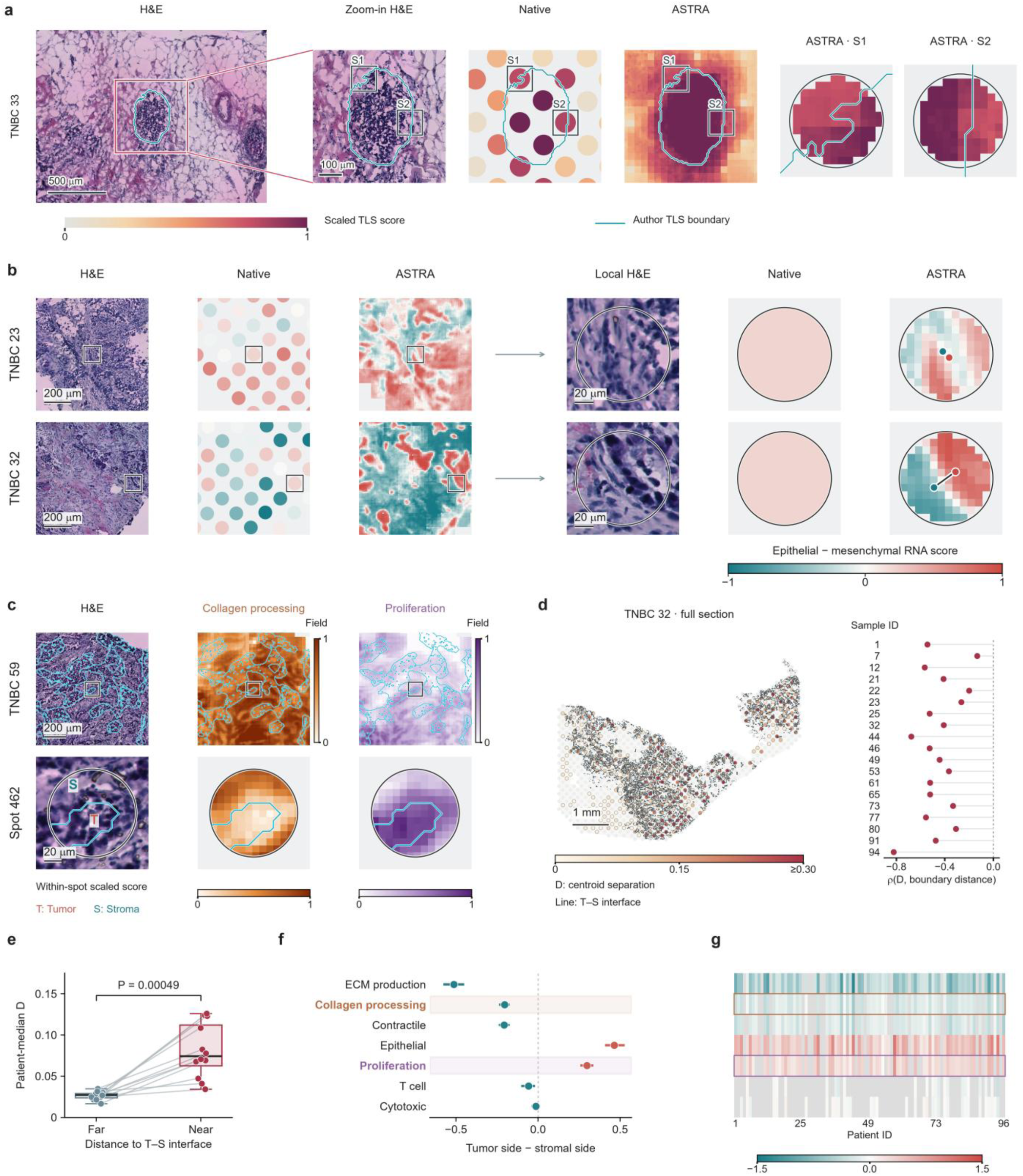
TLS-associated localization and within-spot RNA organization in the TNBC cohort. **a**, H&E, Native and ASTRA TLS-associated scores for an author-annotated TLS in TNBC33. Cyan contours mark the TLS boundary supplied by the original study authors; S1 and S2 denote boundary-crossing spots. **b**, Epithelial–mesenchymal RNA organization in TNBC23/32; colored points mark source centroids. **c**, Collagen-processing and proliferation scores in TNBC59 and spot 462. **d**, Centroid separation D and patient Spearman correlations with tumor–stroma boundary distance (19 patients). **e**, Patient-median D far (≥100 µm) versus near (≤50 µm), 12 patients. Boxes: median/interquartile range; whiskers: 1.5 interquartile ranges; all observations shown. Two-sided exact Wilcoxon P = 0.00049, unadjusted. **f**, Mean tumor-minus-stroma scores and 95% patient-bootstrap intervals (1,999 resamples; n = 76, 73, 69, 76, 68, 22, 35, respectively). **g**, Corresponding values across 92 patients; gray, unavailable. Patient IDs retain original study labels. Scale bars: a, 500/100 µm; b,c, 200/20 µm; d, 1 mm. RNA-source maps and centroids are derived from estimated epithelial and mesenchymal contributions.

Epithelial and mesenchymal RNA-source maps in TNBC23 and TNBC32 showed distinct spatial distributions within individual spots. Collagen-processing and proliferation maps provided molecular context at a tumor–stroma interface in TNBC59 (Fig. 6b,c).

To quantify within-spot organization across the cohort, we defined source separation D as the distance between epithelial- and mesenchymal-RNA-weighted centroids divided by spot diameter. Of the 92 patients, 47 met composite-source support criteria and 20 additionally met reliability criteria for both sources. All 19 patients with evaluable correlations showed greater separation nearer the tumor–stroma interface (Fig. 6d). Among 12 patients with sufficient near (≤50 µm) and far (≥100 µm) spots, patient-median D was higher near the interface in every patient, with a median paired increase of 0.0495 (two-sided exact Wilcoxon P = 0.000488, unadjusted; Fig. 6e).

Program analysis retained the complete 92-patient roster independently of source qualification. Within boundary-crossing spots, ECM-production, collagen-processing and contractile scores favored the stromal side, whereas epithelial and proliferation scores favored the tumor side (Fig. 6f,g). Mean tumor-minus-stroma contrasts were −0.205 for collagen processing (73 patients; 95% interval, −0.231 to −0.180) and +0.297 for proliferation (68 patients; 0.265 to 0.330). Disjoint-gene consistency, measured-HD calibration and a local inference comparison characterized readout reliability (Supplementary Fig. 23). These results extend ASTRA’s cohort-scale application from conventional Visium to the original Spatial Transcriptomics platform, resolving molecular variation within its larger capture spots.

## Discussion

ASTRA combines arbitrary-parent training with frozen inference to make spatial transcriptomic super-resolution reusable across sections and capture formats. In six held-out sections, it achieved higher reconstruction correlations than the evaluated external methods with substantially shorter processing times, and lower errors in the dense-bin benchmark. One source checkpoint supported every primary reconstruction, while optional adaptation further reduced mean dense-bin error and measured-RNA error in the ILC cohort.

The applications connect reconstruction accuracy to spatial interpretation. Controlled CRC experiments showed improved tissue segmentation and tumor-boundary recovery. In real Visium, RNA-source maps and local gene relationships placed a stromal transcriptional state within tissue architecture. CAF and native-HD measurements supported coordinated ECM production and collagen-processing transcription, while ASTRA refined its spatial presentation.

Zero-shot inference then enabled regional and nucleus-associated analysis across the complete 43-section ILC cohort. Arbitrary-parent representation extended the same checkpoint to 92 TNBC sections with larger capture spots, supporting TLS-associated localization and within-spot molecular interpretation. Separating pretraining from deployment makes the same reconstruction model available for cohort analysis across patients and assay formats.

Further validation should extend to additional donors and matched fine-resolution cohort data. Both benchmark geometries use the same six sections, including an adjacent ovarian pair, and scstGCN achieved lower pseudo-Visium nRMSE. Cohort pathology shares H&E with ASTRA’s inputs, while the breast HD calibration reuses a benchmark specimen. The cohort analyses assess anatomical correspondence and consistency of reconstructed RNA patterns. The 8 µm grid specifies output sampling; direct molecular validation at that scale remains an important next step.

The fixed 2,000-gene panel limits molecular coverage. Transcriptomic foundation models such as scGPT and scFoundation^20,21^, and gene embeddings such as GenePT^22^, offer routes to prediction beyond this panel. Incorporating gene-conditioned representations while retaining count conservation and explicit capture geometry could broaden molecular coverage. ASTRA makes pretrained super-resolution practical for analyzing existing spatial transcriptomic datasets and large patient cohorts across assay formats.

## Methods

### Observation representation and expression features

ASTRA receives registered H&E, raw counts, a gene-availability vector and non-overlapping count-bearing regions, termed parents. Parent supports are represented on an internal 2 µm grid; RNA is predicted on an 8 µm grid for a fixed panel of 2,000 genes. For count *C_pg_* of gene g in parent p, with *n_p_* internal cells, the expression input is

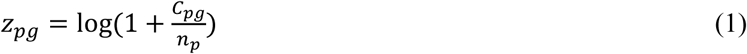

Each support retains the complete region over which its counts were measured. Unavailable genes are masked separately from observed zeros. An assay-conditioned residual adapter handles 3′ expression features, whereas whole-transcriptome probe-based measurements bypass it. A shared encoder combines expression and gene availability. Panel construction, geometry and architectural dimensions are specified in Supplementary Methods 1 and 2 and Supplementary Table 6.

### Morphology and local molecular context

Morphology encoding combines local optical-density and hematoxylin/eosin features, multiscale convolutional features and frozen UNI spatial tokens^13^. Registered features are resampled onto the internal grid without stain normalization. Parent tokens combine expression, support-averaged morphology and geometry. At each output query, attention reads up to eight valid parents ordered by centroid distance, without a fixed radius cutoff; owned cells additionally receive their parent’s token. Morphology, molecular context, coordinates and support indicators are fused into a shared latent representation. Gap cells use an attention-weighted combination of nearby parent densities, with a global fallback when local context is empty (Supplementary Method 2).

### Segment prediction and conservation

An observed segment s is the intersection of a parent support with an output bin; valid unowned cells form a gap segment. A density prior and a segment residual determine the allocation weight

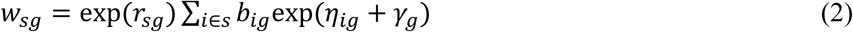

where *b_ig_* is the measured parent density at owned cell i or the local density prior in a gap, *η_ig_* is a bounded spatial residual, *γ_g_* is a bounded gene exposure term and *r_sg_* is a bounded segment residual. For segments *S_p_* belonging to parent p and segments *S_b_* contributing to output bin b,

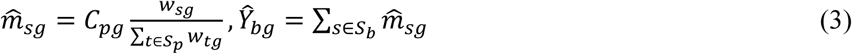

Normalization within each observed parent ensures

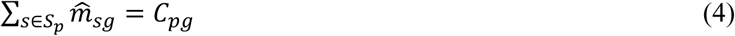

up to floating-point precision. Gap segments retain *m̂*_*sg*_ = *w_sg_* because they have no observed local total. Parent identity is preserved until segment contributions are summed into output bins. Equation (4) applies before averaging overlapping whole-section predictions; this averaging can alter individual parent totals. Numerical implementation and inference settings are detailed in Supplementary Method 2 and Supplementary Table 6.

### Source fitting and optional fine-tuning

The reconstruction network was fitted from scratch on 12 development sections with UNI frozen. Native measurements were aggregated over randomly generated, mutually exclusive parents varying in area, shape and coverage. Supervision used complete, valid 8 µm bins and available genes. The objective was the field-averaged Poisson negative log likelihood (NLL), omitting target-only constants:

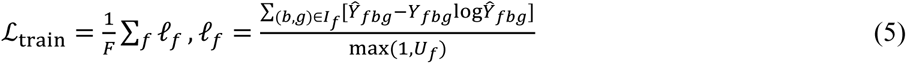

Here F is the number of fields per minibatch, *I_f_* contains scored bin–gene pairs, Y and Ŷ are measured and predicted counts, and *U_f_* is the measured count sum over *I_f_*. The denominator retains zero-count fields as negative examples. Covered, gap and mixed-support bins had equal weights, with no additional parent-level regularizer.

Spatially withheld regions from every development section were evaluated under both HD16 and pseudo-Visium sampling. Checkpoint selection required improvement in at least one task and bounded deterioration in the other, with a fixed initial-score safeguard. Test sections and downstream cohorts contributed neither to panel construction nor source-model selection. Sampling distributions, optimization and stopping rules are specified in Supplementary Method 2.

Optional target fine-tuning used spatially separate coarse-observation training and selection regions. HD16 adaptation reconstructed measured 16 µm counts from 32 µm aggregates. For pseudo-Visium, adjacent spot counts and supports were merged to predict the original spot counts. Coarse Poisson negative log likelihood per UMI was combined with preservation of the frozen model’s allocations:

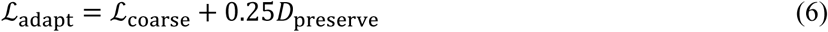

*D*_preserve_ is a count-level Poisson divergence from the frozen source prediction. The reconstruction network was updated while UNI and the source teacher remained frozen. Selection included the initial checkpoint and used only withheld coarse counts; target fine-resolution RNA was excluded from adaptation and selection. Fine-tuning settings are given in Supplementary Method 2 and Supplementary Table 6.

### Expression benchmarking and computation time

FineST, iStar, scstGCN and TESLA were selected to represent contrastive image–RNA learning, hierarchical image features, graph convolution and training-free spatial–histological interpolation^6,8–10^. Pseudo-Visium additionally included SpotZoomer^11^, using one biologically related high-resolution development reference per target, selected before fitting. Native preprocessing, losses and fitting workflows were retained without an additional count-conserving correction. FineST used Virchow2^23^, iStar HIPT^24^, and scstGCN and SpotZoomer UNI^13^. Comparator configurations and reference–target pairs are specified in Supplementary Method 3.

Both benchmarks used the same six held-out sections and 2,000-gene panel. HD16 observations aggregate four aligned 8 µm bins into 16 µm squares; pseudo-Visium observations sum native 2 µm counts within 55 µm circles on a 100 µm lattice. Target 8 µm counts were reserved for evaluation. Frozen ASTRA used one checkpoint; optional ASTRA fine-tuning used coarse target observations. FineST, iStar and scstGCN used target-section fitting, TESLA used interpolation, and SpotZoomer used reference supervision, domain alignment and target fine-tuning.

Gene-wise Pearson correlation coefficient (PCC) was calculated between count-scale predictions and measured counts on common spatial support. Normalized root mean squared error (nRMSE) was calculated after independently min–max scaling prediction and reference for each gene. Position-wise PCC and Bray–Curtis dissimilarity assessed expression profiles across genes. Undefined correlations were omitted, and missing spatial support was not filled. Cross-section summaries equally weighted section means. Gene-wise differences used the highest-PCC external method in each section as the comparator for both metrics and ASTRA variants. Metric-specific eligibility, transformations and display selection are detailed in Supplementary Method 3.

Each run used one NVIDIA A100 GPU (80 GB) and two CPU threads. Runtime summed recorded preparation, target fitting or adaptation, inference and associated processing stages, charging required cache construction once. SpotZoomer totals included reference preparation and reference–target fitting. Pre-existing model pretraining, scoring and figure preparation were excluded. These stage totals are distinguished from cohort queue and accumulated worker times; full timing boundaries are given in Supplementary Methods 3 and 6.

### Colorectal tissue mapping and molecular organization

Pseudo-Visium tissue mapping used a multinomial logistic regression fitted to same-section native expression and molecularly informed pathology labels, then applied unchanged to all reconstructions. MCC, adjusted Rand index, Dice and tumor-boundary metrics were evaluated on common annotated support. Whole-section scores were reported separately from regional averages. *EPCAM* and *COL1A1* profiles used identical pathology-defined distance bands and panel-normalized expression, averaged before log1p transformation. Classifier settings, boundary definitions and regional sensitivities are given in Supplementary Method 4.

For real P2 Visium, a donor-balanced reference excluding P2 supplied RNA-source signatures. Weighted non-negative least squares fitted the same signatures to Native and ASTRA panel-normalized expression, yielding relative RNA contributions. Donor-held-out classification and synthetic mixtures assessed reference performance; program-gene omission tested sensitivity.

Paired gene maps shared Native/ASTRA log1p(CP10K) limits. The joint expression score, J = min(u, v), summarized jointly high relative intensities u and v clipped to 0–1. Disjoint ECM, contractile and collagen-processing gene sets defined stromal transcriptional readouts. Outcomes and covariates were centered within spots; partial correlations adjusted for depth, RNA-source backgrounds, nuclear density and *NOTCH3*. Spatial blocks were the resampling units. Measured non-P2 CAFs, genes outside the prediction panel and native HD expression supplied complementary molecular evidence. Method comparisons characterized an ASTRA-derived candidate. Reference qualification, covariates and gene sets are specified in Supplementary Method 5 and Supplementary Table 4.

### Cohort reconstruction and downstream analysis

The same frozen checkpoint reconstructed all 43 ILC and 92 TNBC sections with explicit input-gene availability. Both registered datasets were excluded from source fitting and panel construction; donor-level exclusion could not be fully audited. Zero-shot refers to reconstruction without updating ASTRA on the target cohort. Quantitative RNA-source and program analyses used directly reconstructed bins fully contained within a unique capture circle (Supplementary Methods 6–8).

ILC regions were selected by pathological class balance, capture coverage, mixed-spot support and boundary requirements, yielding 116 regions from 40 patients. Five-fold patient-held-out classifiers were trained on original-resolution RNA from other patient folds and applied identically to all representations. Regional AUROC, Dice and boundary F1 were averaged equally within patient. Pathology labels were derived from H&E, an input shared with ASTRA. A separate adaptation experiment used five adaptation, three validation and 35 test patients to recover original spot counts from merged observations; adapted weights did not contribute to the zero-shot figures (Supplementary Method 6; Supplementary Table 7).

Donor-balanced breast-cancer single-cell profiles^18^ and weighted non-negative least squares estimated relative RNA contributions. Epithelial scores combined cancer and normal epithelial sources; mesenchymal scores combined CAF and perivascular-like sources. Their difference was evaluated against H&E-derived nuclear contexts within mixed ILC spots. In TNBC, source-centroid separation was related to the tumor–stroma interface. Reference construction, eligibility and measured-HD calibration are specified in Supplementary Methods 7 and 8.

The author-annotated TLS in TNBC33 was selected post hoc for anatomical illustration using a TLS-associated expression score. Seven predefined programs were analyzed across all 92 patients, independently of source qualification. Scores used mean per-gene log1p(CP10K), and tumor-minus-stroma contrasts were aggregated within patient. Gene sets, normalization denominators and spot-level selection rules are given in Supplementary Method 8 and Supplementary Table 4.

### Statistical analysis

Patients were the units of cohort inference. ILC comparisons used two-sided paired Wilcoxon tests; TNBC program contrasts used sign-flip tests. Confidence intervals used patient bootstrapping, and multiple testing used Benjamini–Hochberg correction within the endpoint families defined in Supplementary Methods 6–8^25^. The TNBC near/far comparison used a separate two-sided exact Wilcoxon signed-rank test with unadjusted P. Spatial-block resampling for single-section analyses is specified in Supplementary Methods 4, 5 and 7.

## Manuscript preparation

OpenAI Codex assisted with language editing, manuscript organization, and reference and formatting checks. The authors are responsible for the scientific content and the final text.

## Data availability

All primary measurements originate from the public studies and resources listed in Supplementary Method 1 and Supplementary Table 1. Figure-level tables, eligibility records and data provenance accompany the figures. No new patient material was collected for this computational reanalysis.

## Code availability

ASTRA source code is available at https://github.com/jiachenye-cuilab/ASTRA, with the project website at http://www.cuilab.cn/ASTRA. The accompanying software includes source training, preprocessing, inference and optional target fine-tuning. UNI weights are obtained separately under their original access terms.

## Author contributions

Q.C. designed and supervised the study. J.Y. developed the algorithm, conducted the research and wrote the initial draft. Q.C. revised and edited the manuscript.

